## Supplementary Information for "Modulation of metastable ensemble dynamics explains the inverted-U relationship between tone discriminability and arousal in auditory cortex"

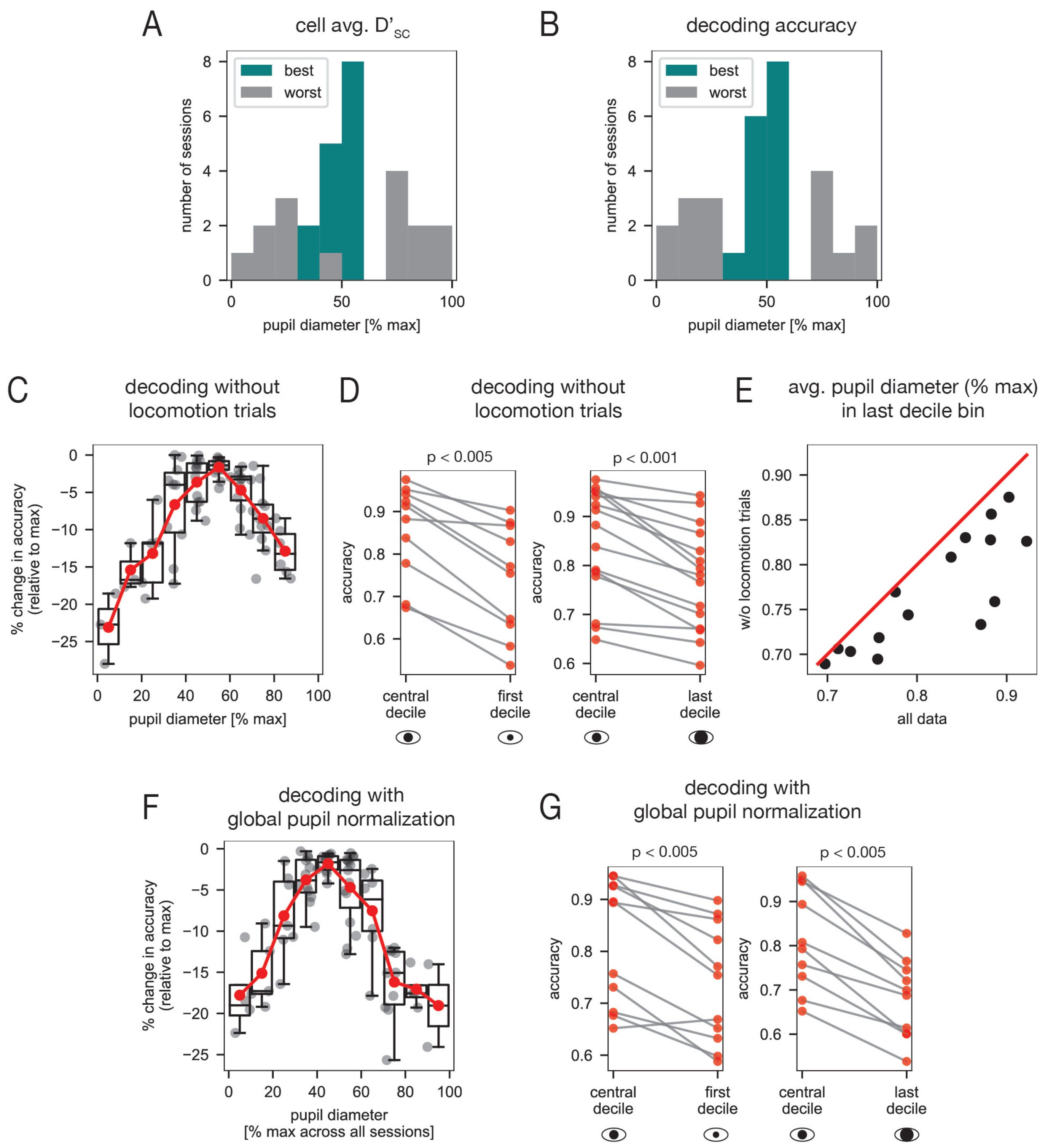


FIG. S1. **Supplementary analyses of stimulus discriminability in the neural data (related to Fig. 2).** **(A,B)** Pupil diameter distributions corresponding to the best and worst cell-averaged $D'_{sc}$ or decoding accuracy. **(A)** In each session, we determined the pupil decile partition for which the cell-averaged $D'_{sc}$ was largest (best decile) or smallest (worst decile). The histogram shows the distribution of the average pupil diameter of the best decile (teal) and worst decile (gray) across all experimental sessions. **(B)** Same as **(A)** but for decoding accuracy.

FIG. S1. (Continued from previous page.) **(C-E)** Session-averaged decoding results when excluding locomotion trials. **(C)** Percent change in cross-validated decoding accuracy (relative session-maximum) *vs.* pupil diameter (group data from 15 sessions). In each pupil diameter bin, we show single-session data (gray) and the corresponding session-average (red) and boxplot. The session-averaged decoding performance still follows an inverted-U with pupil diameter when locomotion trials are discarded. However, without locomotion trials, large pupil diameters are not as robustly expressed (see panel **(E)**) and the right hand side of the inverted-U trend is less distinct compared to the case when all data is used (Fig. 2H). **(D)** *Left*: There is a significant decrease in accuracy in the first pupil decile relative to the most central pupil decile of a session (data from n = 10 sessions with average pupil diameter of first decile $\leq 33\%$ max dilation; $p < 0.005$, Wilcoxon signed-rank test). *Right*: There is a significant decrease in accuracy in the last pupil decile relative to the most central pupil decile of a session (data from n = 15 sessions with average pupil diameter of last decile $\geq 67\%$ max dilation; $p < 0.001$, Wilcoxon signed-rank test). **(E)** The average pupil diameter of trials in the last decile bin of a session without locomotion trials *vs.* when all data is used. The average pupil diameter is noticeably smaller when locomotion trials are excluded. See Methods for methodological details. **(F,G)** Session-averaged decoding results when normalizing the pupil diameter in each session by the global maximum across all sessions, rather than by the maximum within each session separately. **(F)** Percent change in cross-validated decoding accuracy (relative session-maximum) *vs.* (globally-normalized) pupil diameter (group data from 15 sessions). In each pupil diameter bin, we show single-session data (gray) and the corresponding session-average (red) and boxplot. Results are similar to the case of within-session pupil normalization (Fig. 2H). **(G)** *Left*: There is a significant decrease in accuracy in the first pupil decile relative to the most central pupil decile of a session (data from n = 11 sessions with average pupil diameter of first decile $\leq 33\%$ max dilation; $p < 0.005$, Wilcoxon signed-rank test). *Right*: There is a significant decrease in accuracy in the last pupil decile relative to the most central pupil decile of a session (data from n = 10 sessions with average pupil diameter of last decile $\geq 67\%$ max dilation; $p < 0.005$, Wilcoxon signed-rank test). Results are similar to the case of within-session pupil normalization (Fig. 2I). See Methods for methodological details.

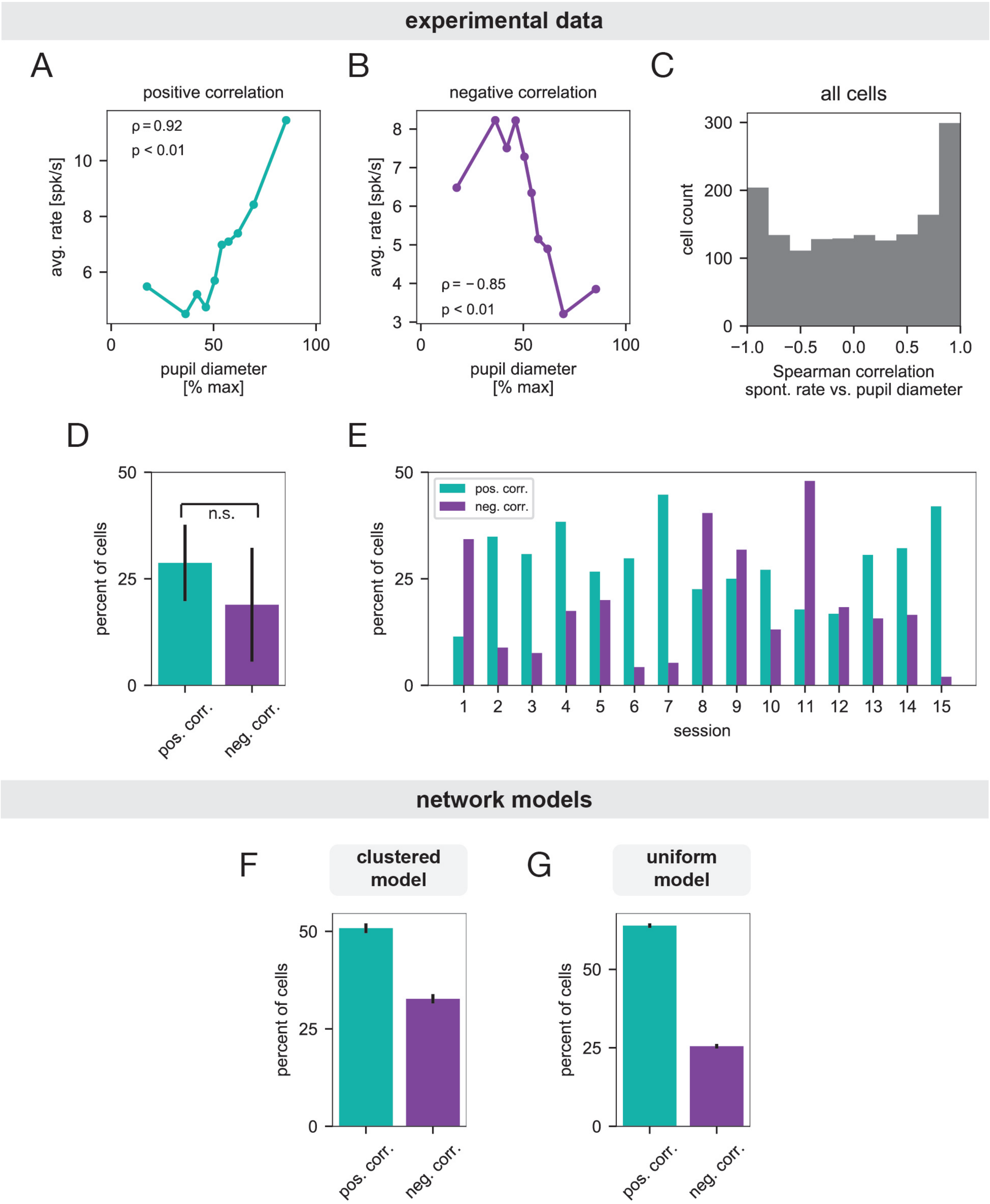


FIG. S2. **Relationships between spontaneous activity and arousal level (related to Fig. 3).**

FIG. S2. (Continued from previous page.) **(A-E) Experimental data.** **(A)** A unit whose spontaneous firing rate increased with pupil diameter (Spearman correlation $\rho = 0.92$, $p < 0.01$). **(B)** A unit whose spontaneous firing rate decreased with pupil diameter (Spearman correlation $\rho = -0.85$, $p < 0.01$. **(C)** Histogram of Spearman correlation coefficients between single-cell spontaneous firing rates and pupil diameter. The histogram includes cells from all experimental sessions. **(D)** Percent of cells whose spontaneous firing rate was significantly positively or negatively correlated with pupil diameter. Bar heights and error bars indicate the mean $\pm$ 1 SD across sessions, and the correlation was considered significant if $p < 0.05$. There was no significant difference between the fraction of positively and negatively modulated units ($p = 0.135$, $n = 15$ sessions, Wilcoxon signed-rank test). **(E)** Percent of cells in each experimental session whose spontaneous firing rate was significantly positively or negatively correlated with pupil diameter. **(F,G) Model networks.** **(F)** Percent of all neurons whose spontaneous firing rate was positively or negatively correlated (Spearman correlation) with arousal level in the clustered network. Bar heights and error bars indicate the mean $\pm$ 1 SD across 10 network realizations, and a correlation was considered significant if $p < 0.05$. **(G)** Same as **(F)** but for the uniform network. The data exhibits heterogeneous dependencies between arousal and ongoing activity, as evidenced by a broad distribution of correlation coefficients between spontaneous firing rate and pupil diameter [panels **(A,B)** for two example units; panel **(C)** for full distribution of correlation coefficients]. Of those units with a significant correlation, there were comparable fractions with positive and negative relationships between ongoing activity levels and pupil-indexed arousal [panel **(D)** for session-average results; panel **(E)** for individual sessions]. Increasing arousal in the network models was also associated with both increases and decreases in spontaneous firing rates [panels **(F)** and **(G)** for clustered model and uniform model, respectively]. The arousal implementation thus qualitatively captures the mixed rate modulations observed in the empirical data. In the clustered model, suppression of excitatory synapses onto pyramidal neurons tends to reduce firing rates, while heightened external drive tends to increase network activity. The diversity of rate modulations thus emerges due to the competition between those two effects, along with the cell-to-cell heterogeneity in the external drive (Fig. 3C). See Methods for methodological details.

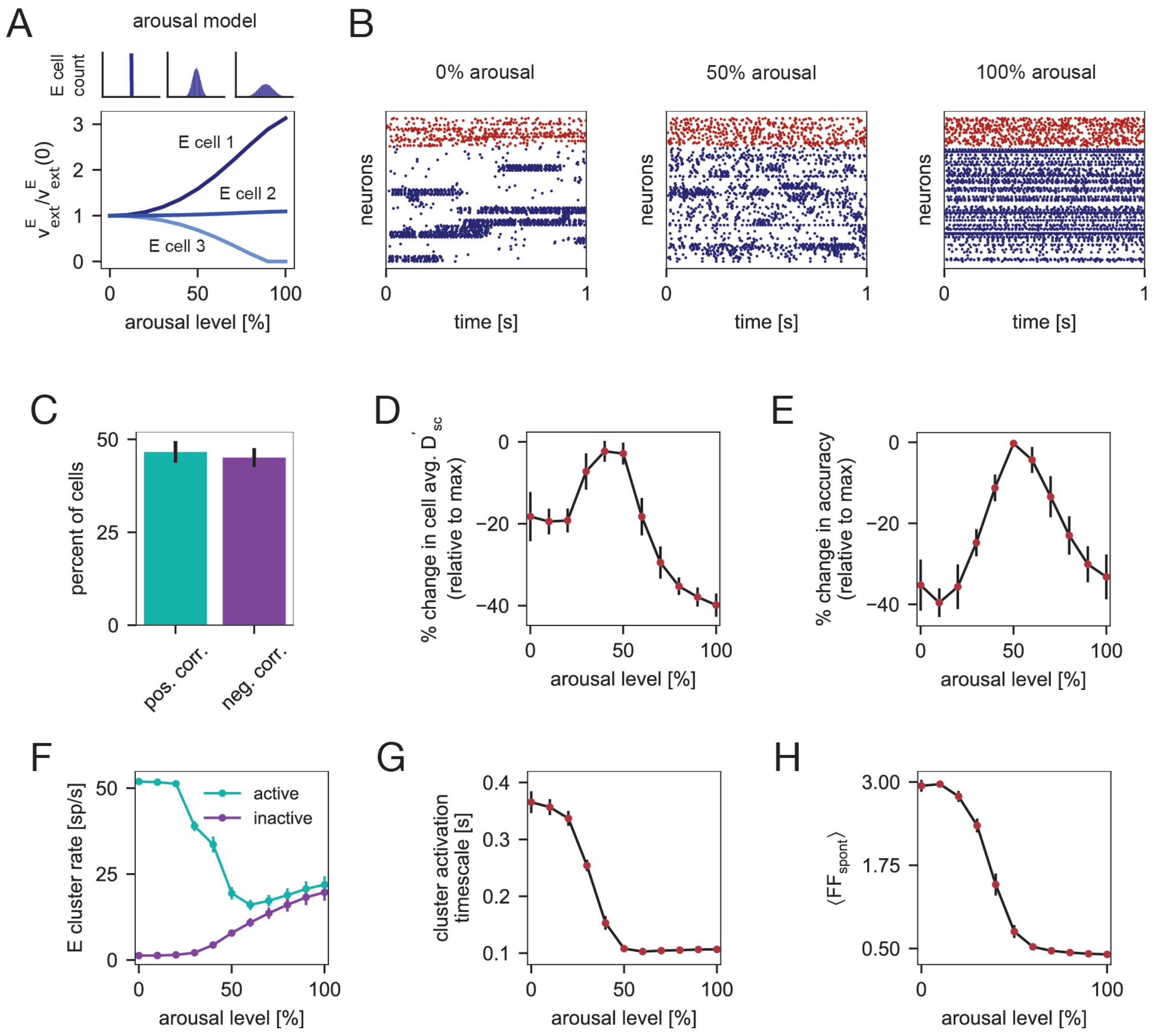


FIG. S3. **A different implementation of arousal induces similar modulations of network activity and stimulus discriminability (related to Figs. 3, 5, 6, and 8).** **(A)** Schematic of an alternative arousal implementation, where arousal is modeled as a heterogeneous modulation of the background inputs to E cells. The lower plot shows the external input $\nu^E_{\text{ext}}$ (relative to its initial value) *vs.* arousal level, where the three different curves correspond to three different excitatory cells. Here, the background input rate to the $i^{th}$ E cell was given by $\nu^E_{\text{ext},i} = \nu^E_o + \Delta^E_{\nu_i}\nu^E_o$, where $\nu^E_o$ is the baseline external input rate to E cells and where $\Delta^E_{\nu_i}$ is a cell-dependent parameter that sets the strength of the modulation to cell $i$. Specifically, $\Delta^E_{\nu_i}$ was given by Eq. 9, with $k = 1.25$, $x_o = 0.275$, $M = 0.9$, and $z^E_i \sim \mathcal{N}(0,1)$ (where $\mathcal{N}(0,1)$ is the standard normal distribution). In this way, increasing arousal increases the variance of the background input rates across cells in the excitatory population, while leaving the spatial average across cells approximately unchanged (inputs were not allowed to go negative). In the clustered model, each assembly was subject to the same realization of the background input distribution, such that all clusters received the same amount of (spatially-averaged) input. **(B)** Example raster plots from simulations of the clustered network at three increasing levels of arousal. **(C)** Percent of all neurons whose spontaneous firing rate was positively or negatively correlated with arousal level in the clustered network (Methods). A mix of positive and negative modulations are observed. **(D)** Percent change in cell-averaged $D'_{sc}$ *vs.* arousal in the clustered model (percent change was computed relative to the maximum across all arousal levels; Methods). The average single-cell discriminability follows an inverted-U relationship with arousal. **(E)** Percent change in cross-validated decoding accuracy *vs.* arousal in the clustered model (percent change was computed relative to the maximum across all arousal levels; Methods). For this analysis, linear classification was performed using population activity from a random sample of 10% of excitatory cells/cluster. The decoding accuracy follows an inverted-U relationship with arousal.

FIG. S3. (Continued from previous page.) **(F)** Average firing rate of active and inactive excitatory clusters as a function of arousal (computed during spontaneous activity). At each arousal level, firing rates correspond to the cluster state with $n_A^*$ active clusters, where $n_A^*$ is the value that occurred most frequently (Methods). The active and inactive cluster rates converge with increasing arousal. **(G)** The average cluster activation timescale *vs.* arousal in simulations of the clustered network (Methods). Cluster activation periods decrease with arousal. **(H)** Population-averaged spontaneous FF ($\mathrm{FF}_{\mathrm{spont}}$) *vs.* arousal (100 ms spike count window; Methods). Neural variability decreases with arousal. Results in this figure are based off of simulations from 5 different network realizations (30 simulated trials/stimulus/network, 5 stimuli; see Methods for simulation details). Data points (circles) and error bars (vertical bars) indicate the mean $\pm$ 1 SD across networks.

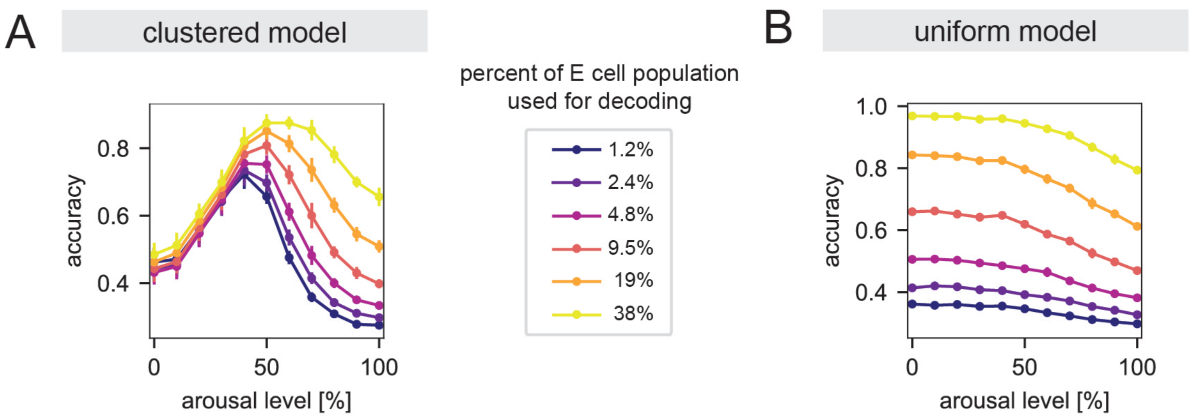


FIG. S4. **Impact of population size on decoding accuracy in the network models (related to Fig. 5).** **(A)** Cross-validated decoding accuracy *vs.* arousal in the clustered model. Different curves show results for different population sizes (displayed as a percent of the total excitatory (E) cell population); data points and error bars indicate the mean $\pm$ 1 SD across network realizations. Note that while decoding accuracy increases with sample size for high arousal, the peak always occurs in an intermediate arousal range for the population sizes considered. **(B)** Same as **(A)** but for the uniform model. For all sample sizes considered, the decoding accuracy decreases with arousal. See Methods for analysis details. In the clustered model, the computational mechanism underlying the inverted-U relationship is a shift in dynamical regime from a metastable attractor phase to a single-attractor uniform phase (Fig. 6). The transition to the uniform phase explains the increase in decoding accuracy with population size in the high arousal regime of the clustered model. Because neurons become independent in the uniform state, adding more neurons averages out variability and improves performance, even though stimulus responses are weak [1]. In terms of population decoding, the inverted-U thus emerges due to competition between response magnitude and response variability. At low arousal, performance is low because variability is high (and pooling has little impact since cells in the same cluster are correlated). At high arousal, a response magnitude-variability tradeoff is present so long as the decoder only samples a subset of neurons from each cluster; then, the decrease in variability obtained by pooling cannot fully compensate for the weak signal, and performance remains low.

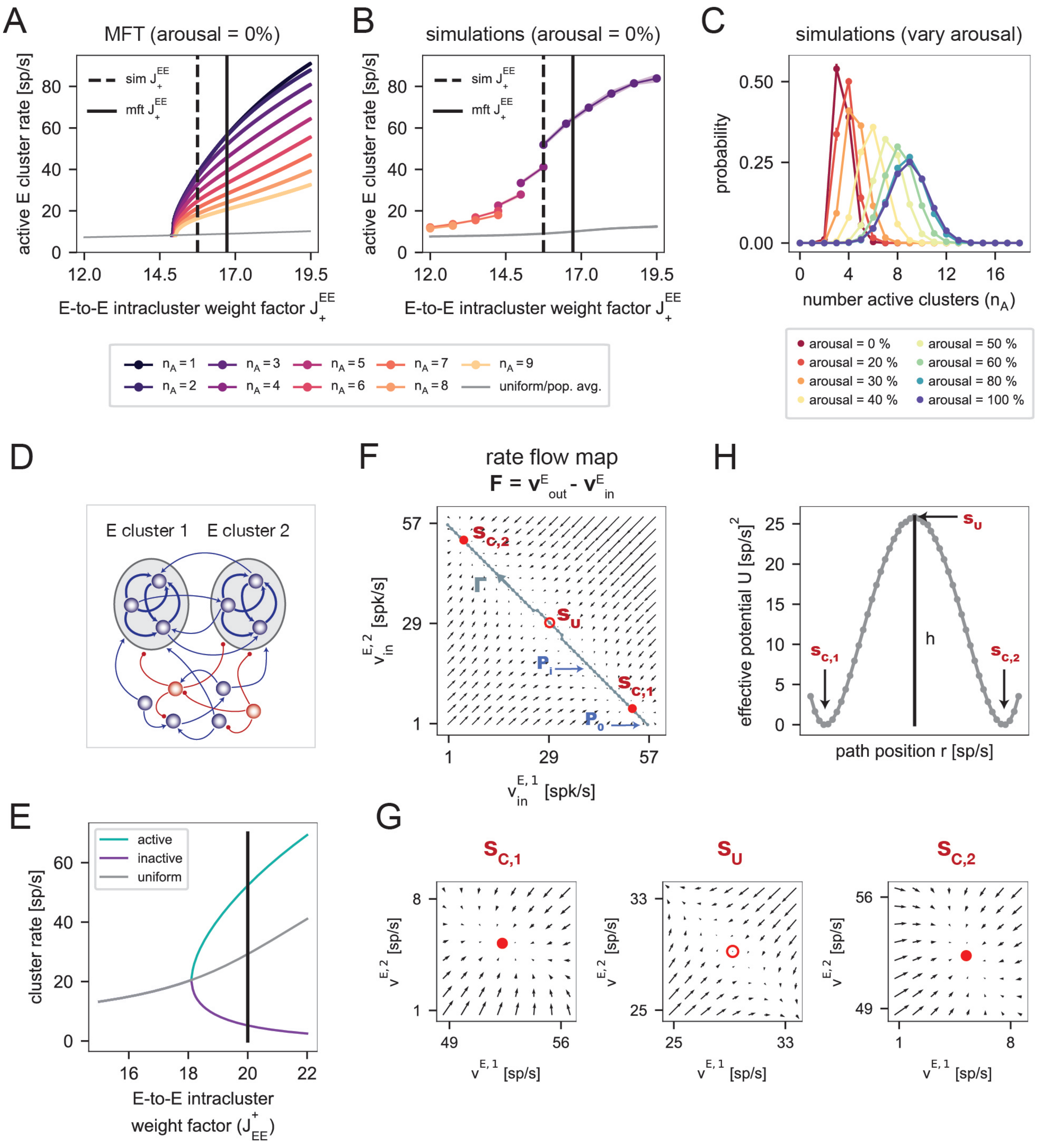


FIG. S5. **Additional details on the mean-field analysis of the clustered model (related to Fig. 6). (A,B)** E-to-E intracluster weight factor controls the onset of cluster states. **(A)** Effect of the E-to-E intracluster weight factor $J_+^{EE}$ on the mean-field solutions of the clustered networks in the absence of arousal. The gray curve shows the rate of the excitatory populations for the solution in which no clusters are active ("uniform" state), and the colored curves show the firing rates of active excitatory clusters for solutions in which $n_A \in \{1, ..., 9\}$ clusters are active ("cluster" states). When $J_+^{EE}$ is below a critical value, the mean-field theory has a single, uniform solution (gray), in which all clusters have the same moderate firing rate. As $J_+^{EE}$ is increased above a critical value, additional solutions emerge. These cluster states are characterized by $n_A \geq 1$ active clusters with a rate $\nu_{n_A,\uparrow}$. Note that the stability of the solutions is not indicated.

FIG. S5. (Continued from previous page.) **(B)** Effect of the E-to-E intracluster weight factor $J_{+}^{EE}$ in the simulations. The gray curve shows the average firing rate of all excitatory neurons and the colored curves show the firing rates of active excitatory clusters conditioned on a particular number $n_A$ of active clusters. At a given $J_{+}^{EE}$, rates are only plotted for values of $n_A$ that occurred with probability $P(n_A) \geq 0.2$. Though there are differences between the theory and simulations (specifically, cluster states emerge at lower $J_{+}^{EE}$ in the simulations), the same qualitative behavior is observed in both cases (the active cluster rate increase with $J_{+}^{EE}$). In both panels, the black dashed line corresponds to the value of the E-to-E weight factor $J_{EE,\mathrm{sim}}^{+}$ that is used in the simulations when studying the impact of arousal, and the black solid line corresponds to the value $J_{EE,\mathrm{mft}}^{+}$ at which the mean-field theory is performed. Note that the arousal-dependent mean-field calculations use a larger $J_{+}^{EE}$ than the simulations ($J_{EE,\mathrm{mft}}^{+} > J_{EE,\mathrm{sim}}^{+}$); the value of $J_{EE,\mathrm{mft}}^{+}$ was chosen to achieve the best match with the simulated firing rates (at $J_{EE,\mathrm{sim}}^{+}$) in the absence of arousal. See Methods for details. **(C)** Probability of observing a certain number of active clusters $n_A$ for different arousal levels in the simulations (Methods). Circular markers and error bars indicate the mean $\pm$ 1 SD across network realizations. **(D-H)** Details on the mean-field analysis of the 2-cluster circuit. **(D)** Schematic of the 2-cluster network, which contains two excitatory clusters and one background excitatory and inhibitory population. **(E)** Effect of the E-to-E intracluster weight factor $J_{EE}^{+}$ on the mean-field solutions of the reduced 2-cluster network (when the arousal level is zero; Methods). When $J_{EE}^{+}$ is below a critical value, the only solution is one in which the two clusters have the same moderate firing rate ("uniform state"). As $J_{EE}^{+}$ is increased above a critical value, an additional solution emerges in which one cluster is active and the other is inactive ("cluster states"), with rates given by the green and purple curves. Note that the stability of the solutions is not indicated. All analyses of the 2-cluster networks in the main text (Fig. 6D,E) were performed at a fixed E-to-E intracluster weight factor of $J_{EE}^{+} = 20$ (black vertical line). **(F)** We studied the dynamics of the 2-cluster network using the effective mean-field theory developed in [2]. To begin, we numerically constructed the rate flow map of the two excitatory clusters, which indicates how the two cluster firing rates will evolve from some initial configuration $\boldsymbol{\nu}_{\mathrm{in}}^{E}$. To accomplish this, we tiled the $\nu_{\mathrm{in}}^{E,1}$-$\nu_{\mathrm{in}}^{E,2}$ plane with a grid, and at each grid location, we computed the induced output rates $\nu_{\mathrm{out}}^{E,1}$ and $\nu_{\mathrm{in}}^{E,2}$ using the effective theory (Methods). Here, the rate flow map is visualized by plotting the vector $\mathbf{F} = \boldsymbol{\nu}_{\mathrm{out}}^{E} - \boldsymbol{\nu}_{\mathrm{in}}^{E}$ at each grid point. From the rate flow diagram, one can identify the three fixed points from the full mean-field theory in **(E)**, corresponding to the uniform solution ($S_U$) and the cluster states in which either the first ($S_{C,1}$) or second ($S_{C,2}$) cluster is active. Moreover, the flow map indicates that the uniform solution is unstable, while the two cluster states are attractors. **(G)** Close-ups of the three fixed points in **(F)**. **(H)** To obtain intuition about transitions between the two attractors, we considered a path $\Gamma$ (gray dotted line in **(F)** connecting the two cluster states $S_{C,1}$ and $S_{C,2}$ through the unstable fixed point $S_U$. For each point $\mathrm{P}_i$ on the path, we computed the line integral $-\int_{\Gamma_{\mathrm{P}_i}} \mathbf{F} \cdot d\boldsymbol{\nu}_{\mathrm{in}}^{E}$, where $\Gamma_{\mathrm{P}_i}$ denotes the segment of the path from $\mathrm{P}_0$ to $\mathrm{P}_i$. This procedure yields a 1-dimensional effective potential $U$, which summarizes the cluster dynamics. Specifically, the potential wells correspond to the two attractors $S_{C,1}$ and $S_{C,2}$, and these configurations are separated by a barrier at the unstable fixed point $S_U$ whose height controls the rate of switching between the two cluster states.

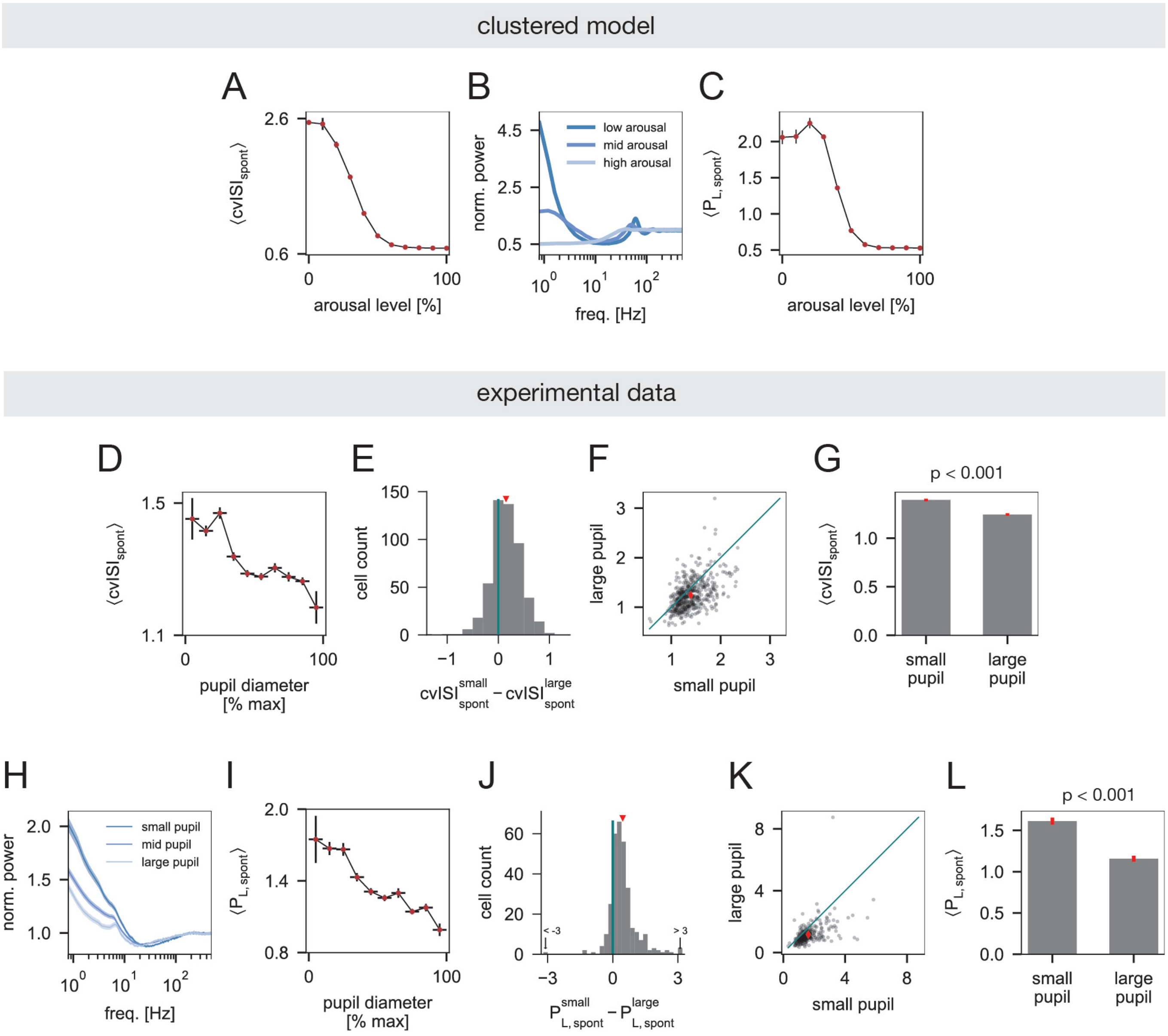


FIG. S6. **Additional measures of spontaneous neural variability in the clustered model and experimental data (related to Fig. 8). (A-C)** Results from the clustered model. **(A)** Population-averaged coefficient of variation of interspike intervals during spontaneous activity ($\mathrm{cvISI_{spont}}$) *vs.* arousal. Data points and error bars indicate the mean $\pm$ 1 SD across network realizations. **(B)** Population-averaged spike-train power spectrum (rate-normalized) of spontaneous activity at low, moderate, and high arousal. **(C)** Same as **(A)**, but for spontaneous low-frequency power (average over 1-4 Hz; $\mathrm{P_{L,spont}}$). **(D-L)** Results from the experimental data. **(D)** Population-averaged $\mathrm{cvISI_{spont}}$ *vs.* pupil diameter (units pooled over sessions). Horizontal error bars indicate pupil diameter bins; data points and vertical error bars indicate mean $\pm$ SEM across cells from all sessions that contribute to the corresponding pupil bin. **(E)** Distribution of the difference in $\mathrm{cvISI_{spont}}$ between small and large pupil diameters (red triangle indicates mean difference). **(F)** $\mathrm{cvISI_{spont}}$ of individual cells in large pupil *vs.* small pupil conditions (red diamond indicates mean values). **(G)** The mean $\pm$ SEM of $\mathrm{cvISI_{spont}}$ in small pupil and large pupil conditions (small pupil: 1.39 $\pm$ 0.01; large pupil: 1.24 $\pm$ 0.01) $\mathrm{cvISI_{spont}}$ is significantly smaller in states of high pupil-indexed arousal compared to low pupil-indexed arousal [$n = 510$ units pooled over 9 sessions with average pupil diameter of smallest (largest) decile bin $\leq 33\%$ ($\geq 67\%$) max dilation; $p < 0.001$, Wilcoxon signed-rank test]. **(H)** Population-averaged spike-train power spectrum (rate-normalized) of spontaneous activity at small ($0 - 33\%$ max dilation), moderate ($33 - 67\%$ max dilation), and large ($67 - 100\%$ max dilation) pupil diameters. **(I)** Same as **(D)** but for population-averaged $\mathrm{P^L_{spont}}$. **(J-L)** Same as **(E-G)** but for $\mathrm{P_{L,spont}}$; $\mathrm{P^L_{spont}}$ is significantly smaller in states of high pupil-indexed arousal compared to low pupil-indexed arousal [small pupil mean $\pm$ SEM: 1.61 $\pm$ 0.04; large pupil mean $\pm$ SEM: 1.16 $\pm$ 0.04; $n = 313$ units pooled over 9 sessions with average pupil diameter of smallest (largest) decile bin $\leq 33\%$ ($\geq 67\%$) max dilation; $p < 0.001$, Wilcoxon signed-rank test]. See Methods for methodological details on the cvISI and power spectra analyses.

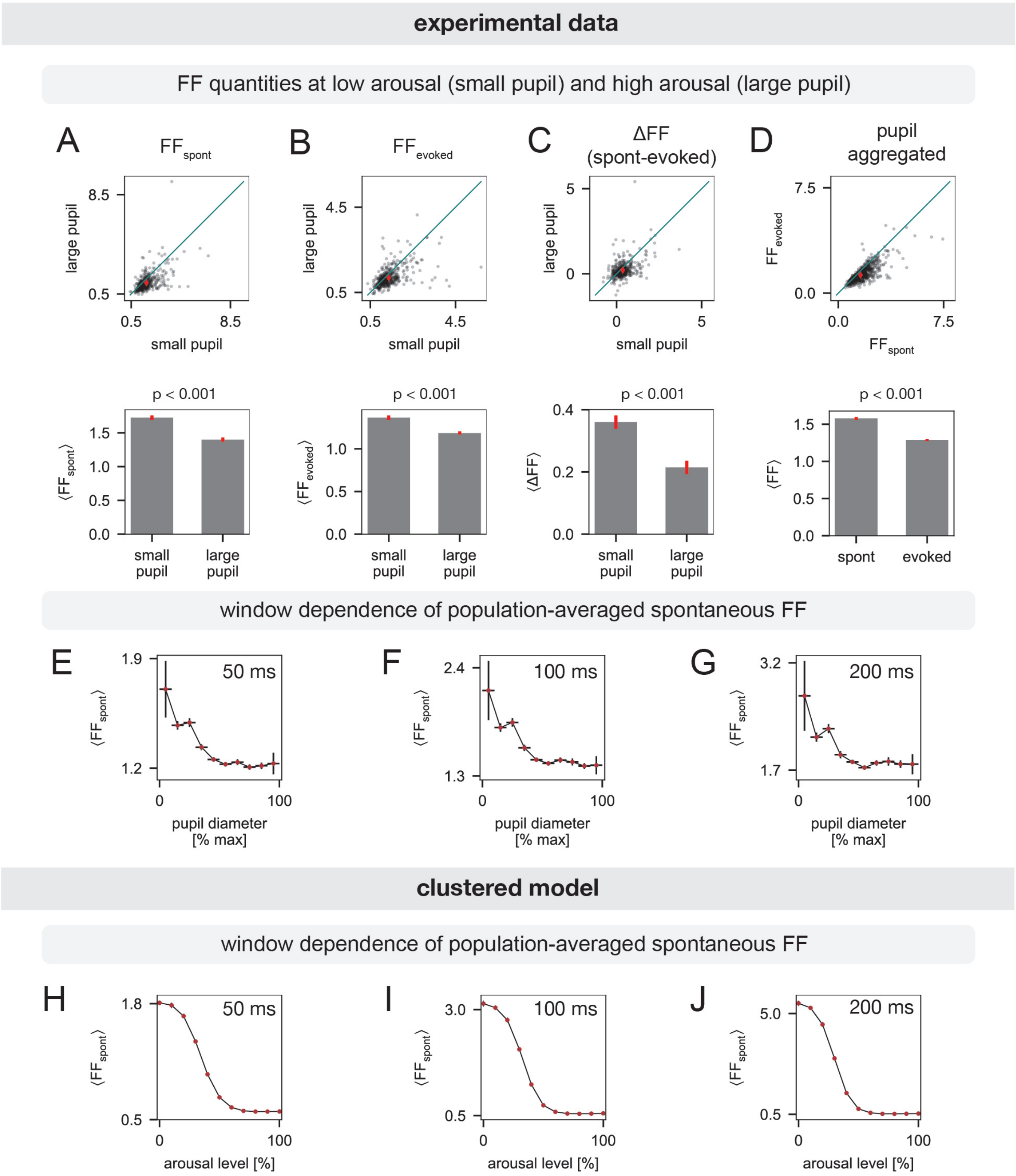


FIG. S7. **Supplementary analyses of the Fano factor (related to Fig. 8). (A-D)** FF quantities at low arousal (small pupil) and high arousal (large) pupil in the experimental data. **(A)** *Top*: Spontaneous Fano factor ($FF_{spont}$) of individual cells in large pupil *vs.* small pupil conditions. The red diamond indicates the mean values. The distribution of single-cell differences is shown in Fig. 8E. *Bottom:* The mean $\pm$ SEM of $FF_{spont}$ in small pupil and large pupil conditions (small pupil: $1.72 \pm 0.03$; large pupil: $1.40 \pm 0.03$). $FF_{spont}$ is significantly smaller in the large pupil condition [$n = 487$ units pooled over 9 sessions with average pupil diameter of smallest (largest) decile bin $\leq 33\%$ ($\geq 67\%$) max dilation; $p < 0.001$, Wilcoxon signed-rank test].

FIG. S7. (Continued from previous page.) **(B)** *Top*: Same as **(A)**, but for $\mathrm{FF}_{\mathrm{evoked}}$. The distribution of single-cell differences is shown in Fig. 8G. *Bottom:* The mean ± SEM of $\mathrm{FF}_{\mathrm{evoked}}$ in small pupil and large pupil conditions (small pupil: 1.36 ± 0.02; large pupil: 1.18 ± 0.02). $\mathrm{FF}_{\mathrm{evoked}}$ is significantly smaller in the large pupil condition ($n = 487$ units, $p < 0.001$, Wilcoxon signed-rank test). **(C)** *Top*: Same as **(A)**, but for $\Delta$FF (spontaneous - evoked). The distribution of single-cell differences is shown in Fig. 8I. *Bottom:* The mean ± SEM of $\Delta$FF in small pupil and large pupil conditions (small pupil: 0.36 ± 0.02; large pupil: 0.21 ± 0.02). $\Delta$FF is significantly smaller in the large pupil condition ($n = 487$ units, $p < 0.001$, Wilcoxon signed-rank test). **(D)** *Top:* Pupil-aggregated $\mathrm{FF}_{\mathrm{evoked}}$ *vs.* $\mathrm{FF}_{\mathrm{spont}}$ of individual cells. The red diamond indicates the mean values. *Bottom:* The mean ± SEM of the pupil-aggregated FF in spontaneous and evoked conditions (spontaneous: 1.58 ± 0.02; evoked: 1.29 ± 0.01). FF is significantly smaller in evoked conditions ($n = 1114$ units pooled over 15 sessions, $p < 0.001$, Wilcoxon signed-rank test). **(E-G)** Population-averaged $\mathrm{FF}_{\mathrm{spont}}$ *vs.* pupil diameter for different window sizes in the experimental data. In each panel, horizontal error bars indicate pupil diameter bins, and data points and vertical error bars indicate the mean ± SEM across cells from all sessions that have sufficient data in the corresponding pupil bin. Spike-count window lengths are indicated in the upper right corner of each plot [panel **(E)** 50 ms, panel **(F)** 100 ms, panel **(G)** 200 ms]. **(H-J)** Population-averaged $\mathrm{FF}_{\mathrm{spont}}$ *vs.* arousal level for different window sizes in the clustered model. In each panel, data points and error bars indicate the mean ± SD of the population-averaged $\mathrm{FF}_{\mathrm{spont}}$ across network realizations. Spike-count window lengths are indicated in the upper right corner of each plot [panel **(E)** 50 ms, panel **(F)** 100 ms, panel **(G)** 200 ms]. In the model, $\mathrm{FF}_{\mathrm{spont}}$ increases with the length of the spike-count window in the low arousal regime, where cluster activity is strong. This is consistent with prior work [3], and indicates the presence of slow rate fluctuations that cause variability to increase over longer integration times. In the data, $\mathrm{FF}_{\mathrm{spont}}$ also increases with window length at low arousal. However, we additionally observe increases in $\mathrm{FF}_{\mathrm{spont}}$ at moderate and high arousal. These effects suggest that the data may have additional sources of variability that are not present in the model and that impact all arousal levels. The low-arousal variation in $\mathrm{FF}_{\mathrm{spont}}$ across window sizes is also less drastic in the data compared to the model, indicating that activity fluctuations are less extreme in the former. Nonetheless, the data exhibits a suppression of neural variability with increasing pupil diameter for all time windows considered. See Methods for methodological details on the FF analyses.

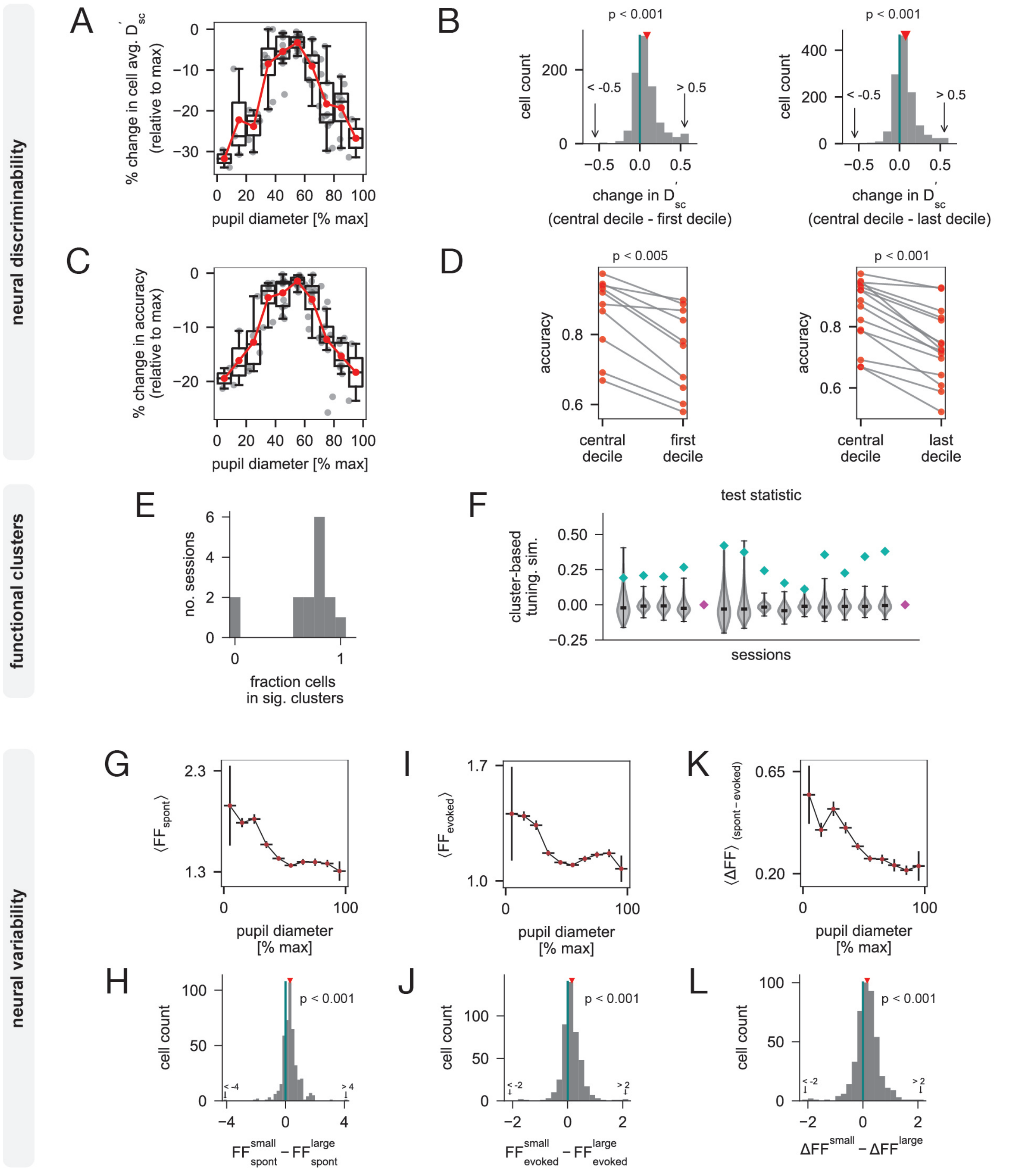


FIG. S8. **Robustness of main results with stricter cell section criteria in the experimental data (related to Figs. 2, 4, and 8).** We examined the robustness of the main results under a more conservative cell selection method (i.e., when implementing strict criteria for sound-responsiveness; see Methods). **(A-D) Neural discriminability analyses** (see Methods for methodological details). **(A)** Percent change in cell-averaged $D'_{\text{sc}}$ (relative to session-maximum) *vs.* pupil diameter (group data from 15 sessions). In each pupil diameter bin, we show single-session data (gray) and the corresponding session-average (red) and boxplot. The population-averaged $D'_{\text{sc}}$ exhibits an inverted-U relationship with pupil diameter.

FIG. S8. (Continued from previous page.) **(B)** *Left:* Difference in $D'_{\rm sc}$ between the most central and first pupil decile of a session (red triangle indicates mean difference). $D'_{\rm sc}$ is significantly smaller in the first pupil decile ($n = 817$ units pooled over 10 sessions with average pupil diameter of first decile $\leq 33\%$ max dilation; $p < 0.001$, Wilcoxon signed-rank test). *Right:* Distribution of the difference in $D'_{\rm sc}$ between the most central and last pupil decile of a session (red triangle indicates mean difference). $D'_{\rm sc}$ is significantly smaller in the last pupil decile ($n = 1211$ units pooled over 15 sessions with average pupil diameter of last decile $\geq 67\%$ max dilation; $p < 0.001$, Wilcoxon signed-rank test). **(C)** Same as **(A)** but for cross-validated decoding accuracy. The average decoding accuracy exhibits an inverted-U relationship with pupil diameter. **(D)** *Left:* Accuracy in the most central pupil decile and the first pupil decile of a session. The accuracy is significantly smaller in the first pupil decile (data from $n = 10$ sessions with average pupil diameter of first decile $\leq 33\%$ max dilation; $p < 0.005$, Wilcoxon signed-rank test). *Right:* Accuracy in the most central pupil decile and the last pupil decile of a session. The accuracy is significantly smaller in the last pupil decile (data from $n = 15$ sessions with average pupil diameter of last decile $\geq 67\%$ max dilation; $p < 0.001$, Wilcoxon signed-rank test). **(E,F) Functional clustering analysis** (see Methods for methodological details). **(E)** Distribution of the fraction of cells in a session that belong to significant correlation-based clusters. Though two sessions do not exhibit significant clusters under the stricter cell selection criteria, the majority of sessions still do. **(F)** Test statistic for cluster-based tuning similarity in each session. Diamonds indicate observed values and violin plots show distributions obtained by permuting cluster labels across cells. Green diamonds indicate a significant result relative to permuted data ($p < 0.05$) and magenta diamonds otherwise. In most sessions, the observed cluster-based tuning similarity significantly exceeds the distribution obtained under the permutation-based null model, suggesting the presence of neural clusters with some functional organization. **(G-L) Neural variability analyses** (see Methods for methodological details). **(G)** Population-averaged $\mathrm{FF}_{\rm spont}$ *vs.* pupil diameter (units pooled over sessions). Horizontal error bars indicate pupil diameter bins; data points and vertical error bars indicate mean $\pm$ SEM across cells from all sessions that contribute to the corresponding pupil bin (note that only one session with relatively few cells contributes to the first pupil bin). The spontaneous FF decreases with pupil diameter. **(H)** Distribution of the difference in $\mathrm{FF}_{\rm spont}$ between small and large pupil diameters (red triangle indicates mean difference). $FF_{\rm spont}$ is significantly smaller in the large pupil condition [$n = 433$ units pooled over 9 sessions with average pupil diameter of smallest (largest) decile bin $\leq 33\%$ ($\geq 67\%$) max dilation; $p < 0.001$, Wilcoxon signed-rank test]. **(I, J)** Same as **(G,H)**, but for $\mathrm{FF}_{\rm evoked}$. Although the decreasing trend in the evoked FF is less drastic with the stricter cell selection, $\mathrm{FF}_{\rm evoked}$ is still significantly smaller for large ($\geq 67\%$ max dilation) pupil diameters relative to small ($\leq 33\%$ max dilation) pupil diameters ($n = 433$ units; $p < 0.001$, Wilcoxon signed-rank test). **(K,L)** Same as **(G,H)**, but for $\Delta$FF (difference between $\mathrm{FF}_{\rm spont}$ and $\mathrm{FF}_{\rm evoked}$). $\Delta$FF generally declines with pupil diameter, and $\Delta$FF is significantly smaller for large ($\geq 67\%$ max dilation) pupil diameters relative to small ($\leq 33\%$ max dilation) pupil diameters ($n = 433$ units; $p < 0.001$, Wilcoxon signed-rank test).

| Parameter | Description | Value |
|---|---|---|
| $N^E$ | number of E cells | 1600 |
| $N^I$ | number of I cells | 400 |
| $\tau_{\mathrm{m}}^E$ | membrane time constant of E cells | 20 ms |
| $\tau_{\mathrm{m}}^I$ | membrane time constant of I cells | 20 ms |
| $\tau_{\mathrm{syn}}^E$ | E synaptic time constant | 5 ms |
| $\tau_{\mathrm{syn}}^I$ | I synaptic time constant | 5 ms |
| $\tau_{\mathrm{ref}}^E$ | refractory period of E cells | 5 ms |
| $\tau_{\mathrm{ref}}^I$ | refractory period of I cells | 5 ms |
| $V_{\mathrm{thresh}}^E$ | threshold potential of E cells | 1.5 mV |
| $V_{\mathrm{thresh}}^I$ | threshold potential of I cells | 0.75 mV |
| $V_{\mathrm{r}}^I$ | reset potential of I cells | 0 mV |
| $V_{\mathrm{r}}^I$ | reset potential of I cells | 0 mV |
| $p^{EE}$ | E-to-E recurrent connectivity fraction | 0.2 |
| $p^{IE}$ | E-to-I recurrent connectivity fraction | 0.5 |
| $p^{EI}$ | I-to-E recurrent connectivity fraction | 0.5 |
| $p^{II}$ | I-to-I recurrent connectivity fraction | 0.5 |
| $J_U^{EE}$ | uniform E-to-E synaptic strength | $0.63/\sqrt{N}$ mV |
| $J_U^{IE}$ | uniform E-to-I synaptic strength | $0.63/\sqrt{N}$ mV |
| $J_U^{EI}$ | uniform I-to-E synaptic strength | $-1.9/\sqrt{N}$ mV |
| $J_U^{II}$ | uniform I-to-I synaptic strength | $-3.8/\sqrt{N}$ mV |
| $p$ | number of E and I clusters | 18 |
| $f^E$ | fraction of E cells/cluster | 0.05 |
| $f^I$ | fraction of I cells/cluster | 0.05 |
| $J_+^{EE}$ | E-to-E intracluster weight factor | 15.75 |
| $J_+^{IE}$ | E-to-I intracluster weight factor | 5.45 |
| $J_+^{EI}$ | I-to-E intracluster weight factor | 6.25 |
| $J_+^{II}$ | I-to-I intracluster weight factor | 5.0 |
| $C_{\mathrm{ext}}^{EE}$ | number of external synapses to E cells | 320 |
| $C_{\mathrm{ext}}^{IE}$ | number of external synapses to I cells | 320 |
| $J_{\mathrm{ext}}^{EE}$ | external E-to-E synaptic strength | $2.3/\sqrt{N}$ mV |
| $J_{\mathrm{ext}}^{IE}$ | external E-to-I synaptic strength | $2.3/\sqrt{N}$ mV |
| $\nu_o^E$ | baseline external input rate to E cells | 7 spks/s |
| $\nu_o^I$ | baseline external input rate to I cells | 7 spks/s |
| $A_{\mathrm{stim}}^E$ | relative stimulus amplitude for E cells | 0.05 |
| $A_{\mathrm{stim}}^I$ | relative stimulus amplitude for I cells | 0 |
| $t_{\mathrm{stim}}$ | stimulus onset time | 1 s |
| $\tau_r$ | stimulus rise time | 75 ms |
| $\tau_d$ | stimulus decay time | 100 ms |
| —— | sampled arousal levels (clustered network) | $[0, 10, 20, 30, 40, 50, 60, 70, 80, 90, 100]\%$ |
| —— | sampled arousal levels (uniform network) | $[0, 10, 20, 30, 40, 50, 60, 70, 80, 90, 100]\%$ |
| * | additional parameters related to the stimulus and arousal implementation are provided in the Methods | |

TABLE S1. **Baseline parameter values for the spiking network model (related to Methods).**

| Parameter | Description | Value |
|---|---|---|
| $N^E$ | number of E cells | 640 |
| $N^I$ | number of I cells | 160 |
| $\tau_{\text{m}}^E$ | membrane time constant of E cells | 20 ms |
| $\tau_{\text{m}}^I$ | membrane time constant of I cells | 20 ms |
| $\tau_{\text{syn}}^E$ | E synaptic time constant | 5 ms |
| $\tau_{\text{syn}}^I$ | I synaptic time constant | 5 ms |
| $\tau_{\text{ref}}^E$ | refractory period of E cells | 5 ms |
| $\tau_{\text{ref}}^I$ | refractory period of I cells | 5 ms |
| $V_{\text{thresh}}^E$ | threshold potential of E cells | 4.86 mV |
| $V_{\text{thresh}}^I$ | threshold potential of I cells | 5.98 mV |
| $V_{\text{r}}^I$ | reset potential of I cells | 0 mV |
| $V_{\text{r}}^I$ | reset potential of I cells | 0 mV |
| $p^{EE}$ | E-to-E recurrent connectivity fraction | 0.2 |
| $p^{IE}$ | E-to-I recurrent connectivity fraction | 0.5 |
| $p^{EI}$ | I-to-E recurrent connectivity fraction | 0.5 |
| $p^{II}$ | I-to-I recurrent connectivity fraction | 0.5 |
| $J_U^{EE}$ | uniform E-to-E synaptic strength | $0.8/\sqrt{N}$ mV |
| $J_U^{IE}$ | uniform E-to-I synaptic strength | $2.5/\sqrt{N}$ mV |
| $J_U^{EI}$ | uniform E-to-I synaptic strength | $-10.6/\sqrt{N}$ mV |
| $J_U^{II}$ | uniform E-to-I synaptic strength | $-9.7/\sqrt{N}$ mV |
| $p$ | number of E and I clusters | 2 |
| $f^E$ | fraction of E cells/cluster | 0.125 |
| $f^I$ | fraction of I cells/cluster | 0 |
| $J_+^{EE}$ | E-to-E intracluster weight factor | 20 |
| $J_+^{IE}$ | E-to-I intracluster weight factor | 1 |
| $J_+^{EI}$ | I-to-E intracluster weight factor | 1 |
| $J_+^{II}$ | I-to-I intracluster weight factor | 1 |
| $C_{\text{ext}}^{EE}$ | number of external synapses to E cells | 128 |
| $C_{\text{ext}}^{IE}$ | number of external synapses to I cells | 128 |
| $J_{\text{ext}}^{EE}$ | external E-to-E synaptic strength | $14.5/\sqrt{N}$ mV |
| $J_{\text{ext}}^{IE}$ | external E-to-I synaptic strength | $12.9/\sqrt{N}$ mV |
| $\nu_o^E$ | baseline external input rate to E cells | 7 spks/s |
| $\nu_o^I$ | baseline external input rate to I cells | 7 spks/s |
| —— | sampled arousal levels for the effective MFT | 50 evenly spaced values in $[0, 100]\%$ |
| * | additional parameters related to the arousal implementation are provided in the Methods | |

TABLE S2. **Baseline parameter values for the reduced 2-cluster network model (related to Methods).**

[S1] Rubén Moreno-Bote, Jeffrey Beck, Ingmar Kanitscheider, Xaq Pitkow, Peter Latham, and Alexandre Pouget. Information-limiting correlations. *Nature neuroscience*, 17(10):1410–1417, 2014.

[S2] Massimo Mascaro and Daniel J Amit. Effective neural response function for collective population states. *Network: Computation in Neural Systems*, 10(4):351–373, 1999.

[S3] Ashok Litwin-Kumar and Brent Doiron. Slow dynamics and high variability in balanced cortical networks with clustered connections. *Nature neuroscience*, 15(11):1498–1505, 2012.
